# HTSplotter: an end-to-end data processing, analysis and visualisation tool for chemical and genetic *in vitro* perturbation screening

**DOI:** 10.1101/2021.09.06.459128

**Authors:** Carolina Nunes, Jasper Anckaert, Fanny De Vloed, Jolien De Wyn, Kaat Durinck, Jo Vandesompele, Frank Speleman, Vanessa Vermeirssen

## Abstract

In biomedical research, high-throughput screening is often applied as it comes with automatization, higher-efficiency, and more and faster results. High-throughput screening experiments encompass drug, drug combination, genetic perturbagen or a combination of genetic and chemical perturbagen screens. These experiments are conducted in real-time assays over time or in an endpoint assay. The data analysis consists of data cleaning and structuring, as well as further data processing and visualisation, which, due to the amount of data, can easily become laborious, time consuming and error-prone. Therefore, several tools have been developed to aid researchers in this process, but these typically focus on specific experimental set-ups and are unable to process data of several time points and genetic-chemical perturbagen screens. To meet these needs, we developed HTSplotter, available as web tool and Python module, which performs automatic data analysis and visualisation of either endpoint or real-time assays from different high-throughput screening experiments: drug, drug combination, genetic perturbagen and genetic-chemical perturbagen screens. HTSplotter implements an algorithm based on conditional statements in order to identify experiment type and controls. After appropriate data normalization, including growth rate normalization, HTSplotter executes downstream analyses such as dose-response relationship and drug synergism assessment by the Bliss independence (BI), Zero Interaction Potency (ZIP) and Highest Single Agent (HAS) methods. All results are exported as a text file and plots are saved in a PDF file. The main advantage of HTSplotter over other available tools is the automatic analysis of genetic-chemical perturbagen screens and real-time assays where growth rate and perturbagen effect results are plotted over time. In conclusion, HTSplotter allows for the automatic end-to-end data processing, analysis and visualisation of various high-throughput *in vitro* cell culture screens, offering major improvements in terms of versatility, efficiency and time over existing tools.

## Introduction

Identification of new and more effective drugs for treatment of diseases such as cancer, bacterial and viral infections, metabolic disorders, and neurodegenerative diseases is a high medical priority (1)(2)(3)(4). Therefore, biomedical researchers are focusing on the search for new druggable targets and combination treatments. In parallel, this presses researchers to pursuit reproducible, effective and efficient experiments leading to more and faster results. In order to address these needs, researchers are implementing experiment automatization, such as robotic cell seeding and liquid dispensing, and/or automatic readouts (2)(5).

Such high-throughput screens typically encompass the evaluation of *in vitro* effects on growth or viability of drugs (single or in combination) and genetic perturbagen screens, such as siRNA and CRISPR screens (6)(7)(8). Additionally, a genetic perturbagen (gene knockdown or overexpression) can be combined with a drug as to evaluate the response in a specific disease context, leading to the identification of certain genes that offer resistance to a set of drugs, or as to identify candidate genes that might work synergistically with drugs (9)(10). The assessment of drug, drug combination and genetic perturbagen screens can be conducted in endpoint or real-time assays. The latter monitors cell proliferation at regular time intervals over days or weeks (e.g. using Incucyte or CytoSPAR machines), while the endpoint assay allows for the assessment of cell viability at single time points (6)(11). Although both assessments can be applied to medium/high-throughput screens, the endpoint assay is the standard option for large HTS scale (6). Regardless of the assay, the cornerstone is to measure the impact of the perturbagen on the cellular response in a specific disease context. Before proceeding to the biological interpretation of the results, the raw data must be processed and analysed. However, the large number of conditions and/or time points generated, by either endpoint or real-time assays, may result in a laborious, time consuming and error-prone data processing and analysis, which may delay and hinder drug development programs (12)(13).

At the very minimum, downstream processing of HTS consists of comparing control and treatment conditions. However, this might not be sufficient to achieve biological interpretation of the data. When a drug is tested in a dosage range, the dose-response relationship is commonly evaluated at a specific time point. Amongst several models used model to fit the dose-response curve, the most common is the four-parameter logistic. It allows the determination of the maximal effect at the highest dose tested (Emax), the dosage at which half of the maximum effect or inhibition is observed (relative IC50/EC50) and the dosage that provokes 50% of the effect or inhibition (absolute IC50/EC50). The relative IC50/EC50 corresponds to the inflection point of the dose-response curve and the absolute IC50/EC50 to the concentration at which half of the response or effect is achieved. The area under the curve (AUC) is an additional metric used to evaluate the drug’s potency, where potency and AUC are inversely proportional (14)(15)(16).

When combining different drugs, one typically aims to quantify the degree of synergism or antagonism. Highest Single Agent (HSA), Loewe additivity, Zero Interaction Potency (ZIP) and Bliss independence (BI) are commonly used models that assume a non-interaction between drugs and compare the observed and expected combination responses (12)(17). These are, however, based on different assumptions. The Loewe model compares the dose-response relationship of individual drugs. HSA compares the effect of drugs in combination to the highest inhibition effect of a single drug, while the BI model compares the combination effect to the expected effect of independent drugs. Lastly, the ZIP model assumes that two non-interacting drugs are expected to incur minimal changes in their dose-response curves upon combination (12)(17)(18)(19). Based on the requirements for each approach, we characterized the Loewe and ZIP as dose-effect models and the HSA and BI as effect-based models (18) (19). Dose-effect based models work if monotherapy dose-response curves are well characterized, which, depending on the response of a cell line to a drug dose range treatment, might not be achievable (19)(20). As for the effect-based models, knowledge of the inhibition effect of its individual components is necessary to predict the combined effect. Hence, effect-based models can be applied in case of combinations of genetic-chemical perturbagens (9). Additionally, the HSA method is less conservative, given that it focuses on the response of a single compound, disregarding effects of competing compounds, increasing the likelihood of detecting false synergism/antagonism.

Seeding density, medium compositions are some of the variables that influence a division rate of cells, which consequently also affect the dose-response curve metrics (IC50, EC50 and AUC) (21). This variability introduce artificial correlations in data, unknown effects of drug action, and introduce unknown complications into biomarker discovery. As an alternative, Hafner *et al*. propose the growth rate (GR) inhibition metrics, as an alternative metrics to characterize a drug response in a real-time assay. The GR metrics have been shown to be robust and decouple any effect that genotype or microenvironment have on division rate from their effect on drug sensitivity. GR metrics are computed by comparing growth rates in the presence and absence of certain perturbations. The GR values between 1 and 0 reflect partial inhibition, 0 denotes cryostasis, and negative values denote cell death (21). In case of drug screens, complementary to the conventional dose-response metric, GR allows the determination of GR_50_ and GR_max_ captures the maximum drug effect on growth rate, which are computed analogouslly to IC50 and Emax, respectively (15)(21). Besides to drug screening, GR can be applied to different experiment types, such as drug combination, genetic and genetic-chemical perturbation assays (21).

In order to facilitate data processing for HTS approaches, dedicated software has been developed to enable high-throughput data analysis, such as BREEZE for drug screens, CellHTS2 for siRNA, and SynergyFinder for drug combinations (12)(21)(22). Depending on the type of experiment, one must organize the data as to fulfil the tool required input structure. Since these tools are designed for endpoint assays, data resulting from real-time assays implies repetitive work from the user. Moreover, these tools are not suited to analyse combinations of chemical and genetic perturbagens, requiring researcher to conduct their own data analysis manually. To the best of our knowledge, there is currently no software with the ability of implementing automatic end-to-end analyses irrespective of HTS experiment and irrespective of assay type, endpoint or real-time.

To address these limitations, we developed HTSplotter as a freely available web tool and Python module. HTSplotter is tailored to analyse drug, drug combination, genetic perturbagen and combinations of genetic-chemical perturbagen screens, either in real-time or as endpoint. HTSplotter identifies the type of experimental setup through a conditional statement algorithm. It performs a normalization, growth rate profile and, in case of a drug screen, drug combination or genetic-chemical perturbagen experiment, identifies the dose-response relationship for each drug alone. Synergism/antagonism of drug or genetic-chemical combination screens is determined through the BI, HAS or ZIP methods. For a real-time assay, regardless the experiment type of screens, HTSplotter determines the GR of each perturbation in relation to the control, showing the results in a GR plot over time. Finally, results are plotted and exported as PDF files, allowing a fast biological interpretation of the data and efficient end-to-end analyses, irrespectively of either experiment type and assay.

## Results

### HTSplotter: an end-to-end tool for HTS

HTSplotter is freely available as a web-application, https://htsplotter.cmgg.be/, and as a Python library, https://github.com/CBIGR/HTSplotter, together with an outlined documentation including step-by-step user instructions and examples. This tool is written in Python 3.9 using SciPy, h5py, NumPy, Matplotlib, math, os and sys libraries. It automatically identifies four experiment types: drug, drug combination, genetic perturbagen and genetic-chemical perturbagen screens. A genetic perturbagen screen can be a simple knockdown or overexpression of a gene, a CRISPR/Cas9 screen or a siRNA library, while the genetic-chemical perturbagen screen consist of a genetic perturbagen in combination with a drug. These can be conducted either in endpoint or real-time assays.

The evaluation of each experiment type consists of a normalization relative to the control. Additionally, in case of drug, drug combination and genetic-chemical perturbagen screens, the dose-response relationship is calculated for each drug. To determine the synergism/antagonism of a drug combination, the user can choose between the HSA, ZIP and BI method. For genetic-chemical perturbagen screens, the user can choose between the HAS and BI method. The combinations can be higher order or pairwise and therefore there is no limit on the number of combined drugs or the dosage tested. Combinations can be arranged in a matrix *A*, with *N* dimensions and 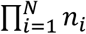 entries, where *N* is the number of drugs, *n*_*i*_ is the number of concentrations of drug *i*. Additionally, for all experiment types, HTSplotter has no limits regarding the number of cell lines tested on each experiment. Fig 1 summarizes the analyses and visualisations provided by HTSplotter.

**Fig 1:**
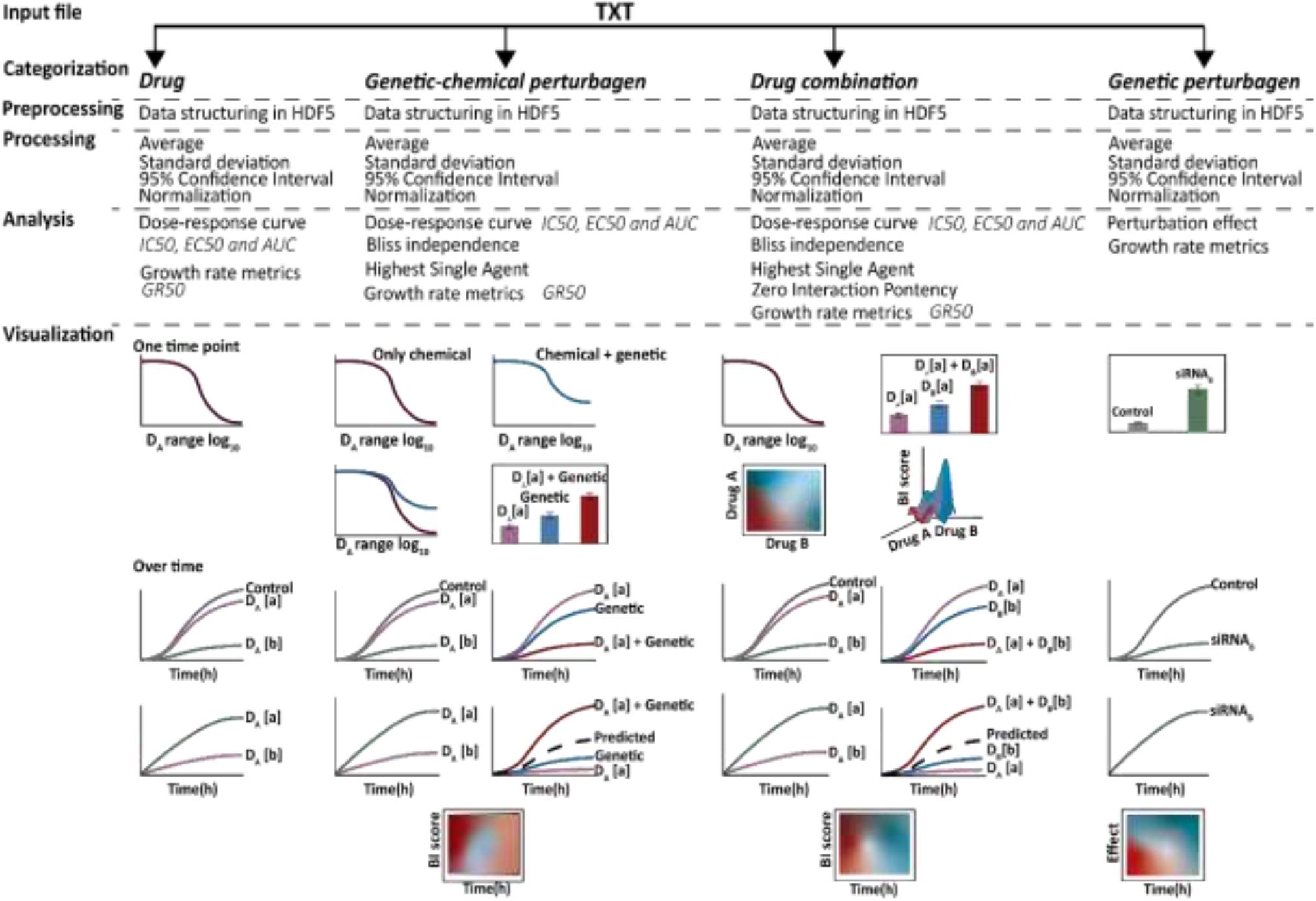
Overview of HTSplotter steps for each type of HTS experiment. The input file, directly imported from HTS machines like Incucyte S3 as a TXT file, is automatically processed and analysed by HTSplotter. As output, TXT and PDF files are generated. The PDF file contains the plots from each type of analysis. In case of a real-time assay, a growth rate is plotted over time. In case of a drug screen a dose-response curve with the growth rate data across the concentrations tested for a time point is depicted. ***Genetic perturbagen screen:*** a heatmap and bar plots are provided with all the perturbagens tested, for specific time points. Additionally, a XY-plot and growth rate plots are created for each perturbagen. ***Drug screen***: a dose-response curve of each drug alone is plotted with associated statistical parameters. XY-plots are also returned with all concentrations of each tested drug over time. Additionally, growth rate plots are provided over time as well as dose-response plots from the growth rate metrics, such as GR50. ***Drug combinations and genetic-chemical perturbagen screens***: a dose-response curve of each drug alone is provided with their statistical parameters. In case of genetic-chemical perturbagen, as to allow for a direct comparison of drug response without and with the perturbagen, a dose-response curve for both situations is plotted. Also, for specific time points, a bar plot containing information about each condition is given. Moreover, XY-plots and growth rate plots, visualize all concentrations of each tested drug over time as well as each combination with respective drugs alone. A heatmap with synergism/antagonism phenotypes scores over time is created, for either drug combination or genetic-chemical combination screens. Additionally, in case of a drug combination, resulting in a combination matrix of two drug (*m* × *n, where m* ≥ 2), a heatmap for each main time point (2D and 3D) is given, whilst in case of *m* = 1 only a 2D heatmap is plotted. HDF5: hierarchical data format file; relative or absolute IC50: 50 % of inhibitory concentration; relative or absolute EC50: effective concentration; AUC: area under the curve; GR_50_: 50 % of growth rate; GR_max_: maximum drug effect.

### HTSplotter: from input file to analysis

HTSplotter utilizes text files directly exported from real-time devices, such as Incucyte and xCELLigence, as inputs, where comment lines above the data to be processed are allowed. “Date Time” and “Elapsed” strings must be present at the first and second data columns respectively, so that the software detects the data headers row (Fig 2**Error! Reference source not found**.), holding the description of the experimental conditions. These columns have date and time points corresponding to each measurement, respectively, e.g. apoptosis, confluency or impedance. In case of more than one time point, defined in this paper as time point intervals, the measurements correspond to time increments from the start of the experiment e.g. 2h, 12h, etc. The information regarding each experimental condition should contain drug name, gene knockdown or overexpression, concentration, units, cell line name and seeding density in one column separated by commas, thus “.” should be used as decimal separator. More detailed information regarding data and header structure are available at the HTSplotter website, as well as data examples for each type of experiment (https://htsplotter.cmgg.be/).

**Fig 2:**
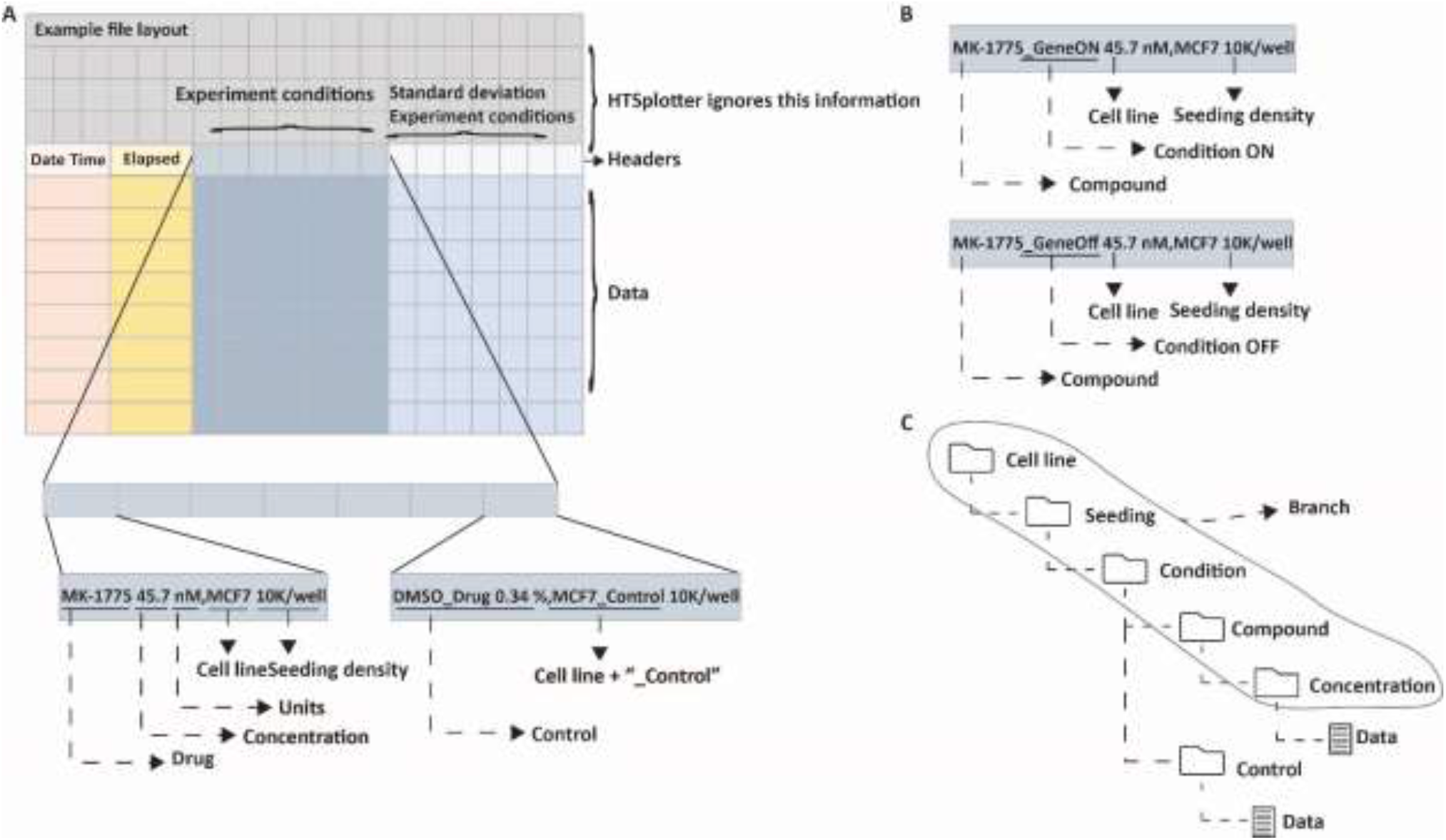
An example input file structure, experimental condition and data structure. **A)** Example of an Input file structure with a detailed view on the experiment condition description. Header and data consist of four sub-groups: “Date Time”, “Elapsed”, “Experiment”. On the left is the description of the MK-1775 drug at a dosage of 45.7 nM on the MCF-7 cell line with 10k cell seeding per well and on the right the control condition, DMSO at 0.34% on the MCF-7 cell line with 10K seeding per well. **B)** Example of an experimental condition description for a genetic-chemical perturbagen. **C)** Data storage structure in the HDF5 file. A branch is created when cell line, seeding, condition and drug are grouped in the same experimental set-up.

HTSplotter implements an algorithm based on conditional statements that extracts information about the number of cell lines, seeding density, conditions, drugs and concentrations tested. In this way, information about similar experiments in a cell line with the same seeding density and conditions for multiple tested drugs, are grouped creating a branch. The conditions can also be gene knockdown or overexpression. The experiment is then categorized as drug, drug combination, genetic perturbagen or genetic-chemical perturbagen screen according to the number of branches, number of identified controls and number of concentrations tested per drug. Categorization is a crucial step for HTSplotter, as it leads to a proper evaluation of the perturbagen of each condition and to the implementation of downstream analyses required for each experiment type, therefore requiring user confirmation about predicted categorization and identified control(s). In case of incorrect categorization, an error file with information regarding data headers and possible missing information is created. Otherwise, the experiment information is stored in a hierarchical data format file (HDF5), the structure of which is determined by the type of experiment. The HDF5 file includes all information from the independent experiments.

Subsequently, for each time point HTSplotter normalizes the data to the proper control(s). The resulting phenotypic measurement (e.g. confluency or apoptosis) can be inhibition or enhancement, this option being given to the user before the analysis. If the standard deviation is present in the input file, the tool rescales it, by dividing the standard deviation of a condition by the standard deviation of the proper control.

In case of repetitive conditions, being these technical or biological replicates in the same file or in independent files, HTSplotter computes averages, standard deviations and 95% confidence intervals. For genetic perturbagen screens with more than one control, HTSplotter calculates averages, standard deviations and 95% confidence intervals of all controls as to have a unique control, and only then the normalization takes place.

Upon a minimum of two concentrations in a drug, drug combination or genetic-chemical perturbagen screen, a dose-response curve of each tested drug and synergism/antagonism using BI, HAS and ZIP methods are calculated (Materials and Methods). Upon drug response in a real-time assay, GR metrics are also calculated (Materials and Methods).

### HTSplotter: visualisation

HTSplotter allows the analyses of endpoint and real-time assays, regardless of the experiment type. If the input file contains more than one time point, additional plots with time features are provided, as well as plots at preselected time points (e.g. 24h, 48h, etc.). A feature of HTSplotter is the XY-plot, where the read-out (e.g. confluency or apoptosis) and either inhibition or enhancement effects (Y) are plotted over time (X) leading to better biological interpretations of the chemical or genetic perturbation effect. Upon drug or genetic-chemical combinations, the predicted combination effect is plotted together with the observed combined and single effect. In this way, XY-plots and bar-plots allow a detailed observation of the synergism/antagonism scores. Additionally, for only real-time assays, regardless the experiment type, HTSplotter provides GR plots, where a growth rate over time for each condition can be visualized. Another unique feature of HTSplotter is the visualisation of synergism scores over time in a heatmap, allowing to identify the onset of synergism/antagonism. In case of drug combinations, when the number of concentrations is higher or equal to two, 2D and 3D heatmaps are shown for main time points. Whilst in the case of a single concentration, only a 2D heatmap is shown. For all HTS drugging experiments, dose-response curves are displayed for main time points, for the experiments where at least two concentrations of a drug were tested.

### HTSplotter analysis of a continuous *in vitro* cell based genetic-chemical perturbagen screen: a case study

Genetic-chemical perturbagen experiments provide phenotypically information on the compound mechanism in combination with a genetic perturbation, resulting in insights on resistance or increased sensitivity phenotype, as well as a synergistic/antagonist combination. Additionally, this experiment type allows the identification of biomarkers for certain treatments and the discovery of novel targets that could work in combination in a certain disease.

In order to illustrate the advantage of HTSplotter on genetic-chemical perturbagen assessed in real-time, we applied our tool to an in-house assessment of exogenous overexpression of the SOX11 gene in the SH-EP neuroblastoma cell line (23), in combination with the MYB inhibitor celastrol (24). MYB protein levels were previously shown to be strongly induced upon SOX11 overexpression (SOX11 OE) (23), while celastrol has been reported to inhibit MYB activity in cancer cells (acute myeloid leukemia) (24)(25)(26). We evaluated the effect of a celastrol dose-range over time in the SH-EP cell line with SOX11 OE versus parental cells. The data used in HTSplotter was directly exported from Incucyte S3, and consisted of 18 different conditions and 36 time points, resulting in 450 data points. The cell line response upon SOX11 OE was assessed by a dose-response, which provided the conventional metrics (abs. IC50, rel. IC50) as well as metrics based on the normalized growth rate inhibition (GR_abs. 50_, GR_rel 50_). The latter are computed by comparing growth rates in the presence and absence of drugs and normalized by the cells doubling time. From both dose-response curves, at 72h, the tested dose-range of celastrol in the induced SOX11 OE (Fig 3**Error! Reference source not found**.), showed lower absolute IC50 and GR_IC50_ values (387.45 nM and 322.12 nM, respectively), when compared to the parental cell line (659.69 nM, 592.58 nM, respectively), indicating increased sensitivity followingSOX11 OE. Additionally, comparing the conventional and GR metrics from 24h, 48h and 72h (S Fig 2), we observed a stabilization on the GR metrics from the 48h, while the conventional ones show more variability. The GR metrics characterizes the drug regardless of time point, conferring a more robust metric when compared with the conventional ones, as these have a higher variability regardless the time point of the analysis, as shown by Hafner *et al*.(21).

**Fig 3:**
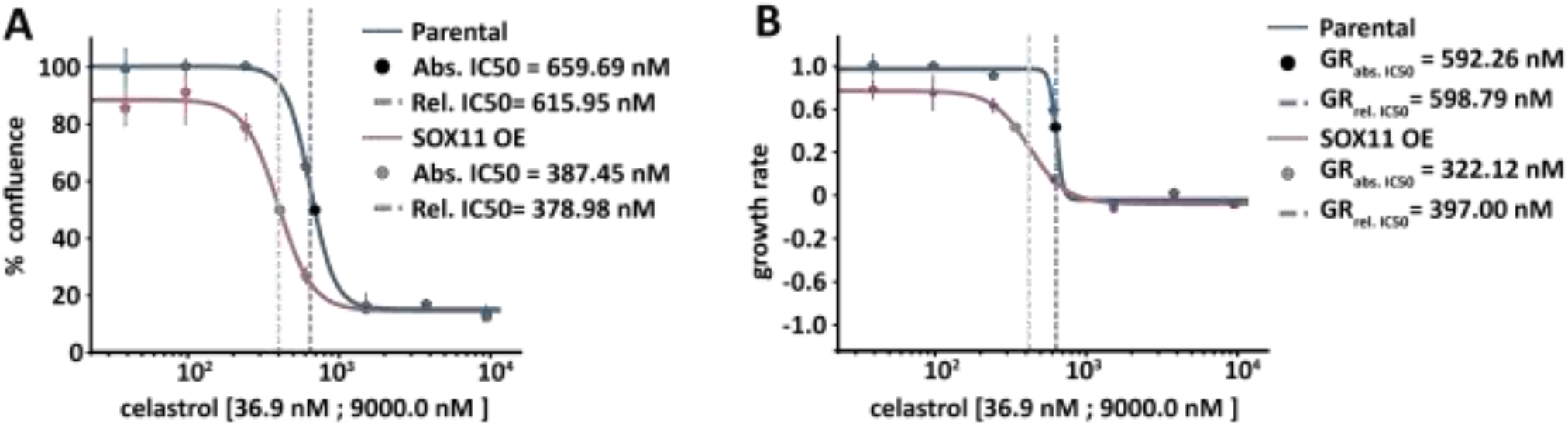
Dose-response curve provided by HTSplotter for induced overexpression of SOX11 in comparison to the parental cell line. **A)** Upon SOX11 OE, the absolute (abs.) and relative (rel.) IC50 shifted to a lower dose (387.45 nM, 378.98 nM, respectively), when compared to the parental cell line (659.69 nM, 615.95 nM, respectively). **B)** GR dose-response curve, upon SOX11 OE, the GR absolute and relative IC50 shifted to a lower dose (322.12 nM, 397 nM, respectively), when compared to the parental cell line (592.58 nM, 599.24 nM, respectively). SOX11 OE: SOX11 overexpression.

In order to identify synergistic phenotypic responses between the gene and drug combinations, HTSplotter applied the BI method, which is more conservative than HAS method. The BI scores are shown over time in the heatmap (Fig 4), where an increasing synergistic effect can be observed between SOX11 OE and celastrol at 576.0 nM, starting at 10h of treatment.

**Fig 4:**
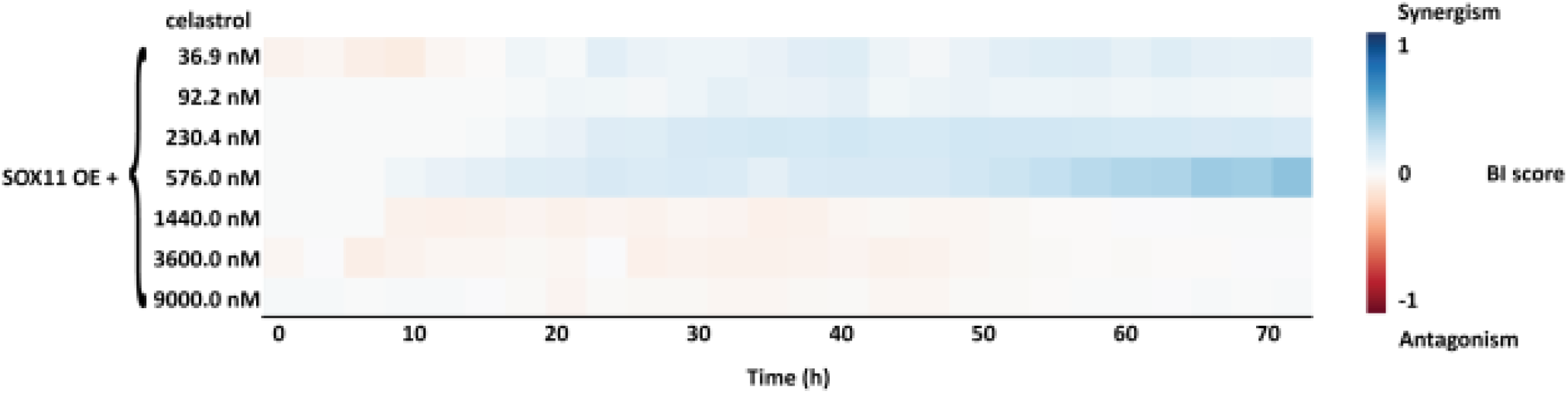
BI score heatmap of effect upon SOX11 overexpression and celastrol dose-range over time. The blue gradient indicates synergism and the red gradient indicates the antagonism between SOX11 overexpression (OE) and celastrol.

Assessing in detail the highest BI score of the genetic-chemical combination (Fig 5, A), celastrol only treated cells showed a higher inhibitory effect than SOX11 OE condition alone. However, after 50h of treatment the inhibitory effect decreased. In contrast, the combination of both genetic-chemical perturbation resulted in an increasing inhibition over time until 50h after treatment, becoming constant until 72h. HTSplotter also allows to plot in terms of growth rate for each condition related to the control (Fig 5, B). Each condition alone (respectively SOX11 OE and celastrol) had a lower but still positive growth rate, while the combined nearly complete growth arrest was noted when compared with the control.

**Fig 5:**
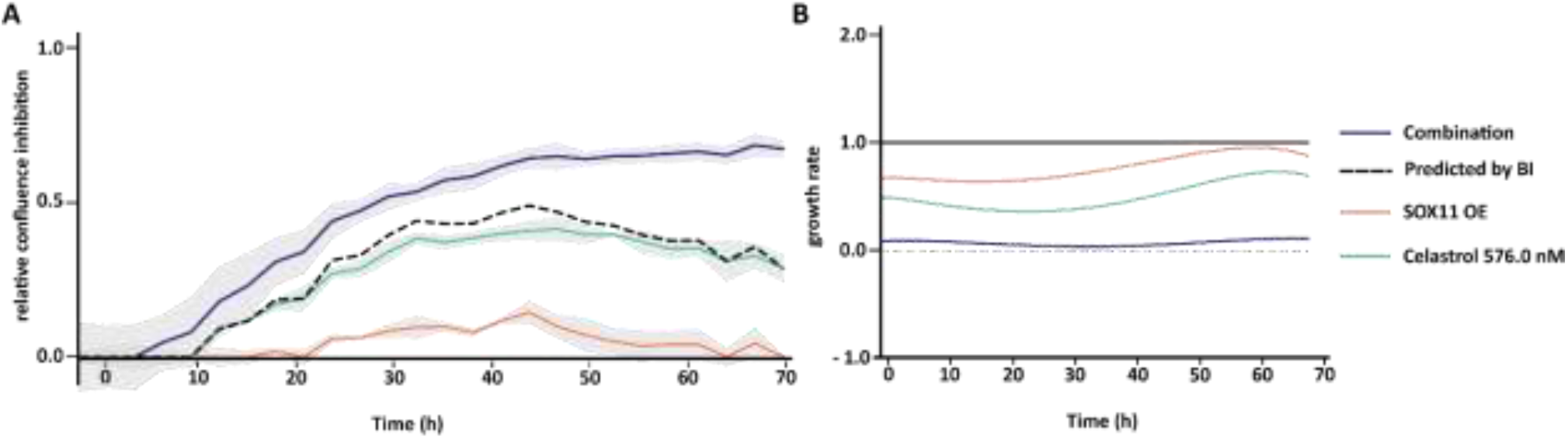
XY-plot and growth rate plot from the genetic-chemical perturbation, SOX11 overexpression (OE) and celastrol. **A)** Relative confluence is the confluence relative to the control. The dash line indicates the predicted combination effect, computed according to the BI method, equation (4). Relative confluence inhibition over time of celastrol at 576.0 nM combined with SOX11 OE. The BI scores at 24h, 48h and 72h are 0.16, 0.16 and 0.35, respectively. **B)** The growth is computed according to equation (7). Growth halt is indicated by the grey dash line. The SOX11 OE and celastrol conditions alone have growth rates lower than the control. The combination of both however, resulted in a almost halted growth rate. SOX11 OE growth rates at 24h, 48h and 72h are 0.65, 0.86, 0.79, respectively, the celastrol (576.0 nM) growth rates at 24h, 48h and 72h are 0.37, 0.55, 0.63, respectively, and the combination growth rates at 24h, 48h and 72h are 0.06, 0.06, 0.1, respectively.

Together, the growing inhibition effect and the growth rate results, observed for the combined condition, suggest an impediment of cell proliferation, while in single conditions the cells are still able to proliferate, although with lower growth rates. Here we show that in a high-throughput manner HTSplotter provides an efficient analysis of the data for genetic-chemical perturbagen experiment, allowing the researcher to focus on the interpretation of the results and answer biological questions.

### HTSplotter analysis of a continuous assessment of *in vitro* cell based drug combinations: a case study

Drug combinations require multiple conditions which, when monitored over time, rapidly multiplies the number of data points thus increasing the number of needed computations. HTSplotter organizes the data according to the experimental set-up to easily implement further data processing, analysis and visualisation. Data processing consists of a normalization of all conditions to the proper control. Analyses include the determination of the dose-response relationship (Fig 6) and synergism/antagonism (Fig 7 and Fig 8).

**Fig 6:**
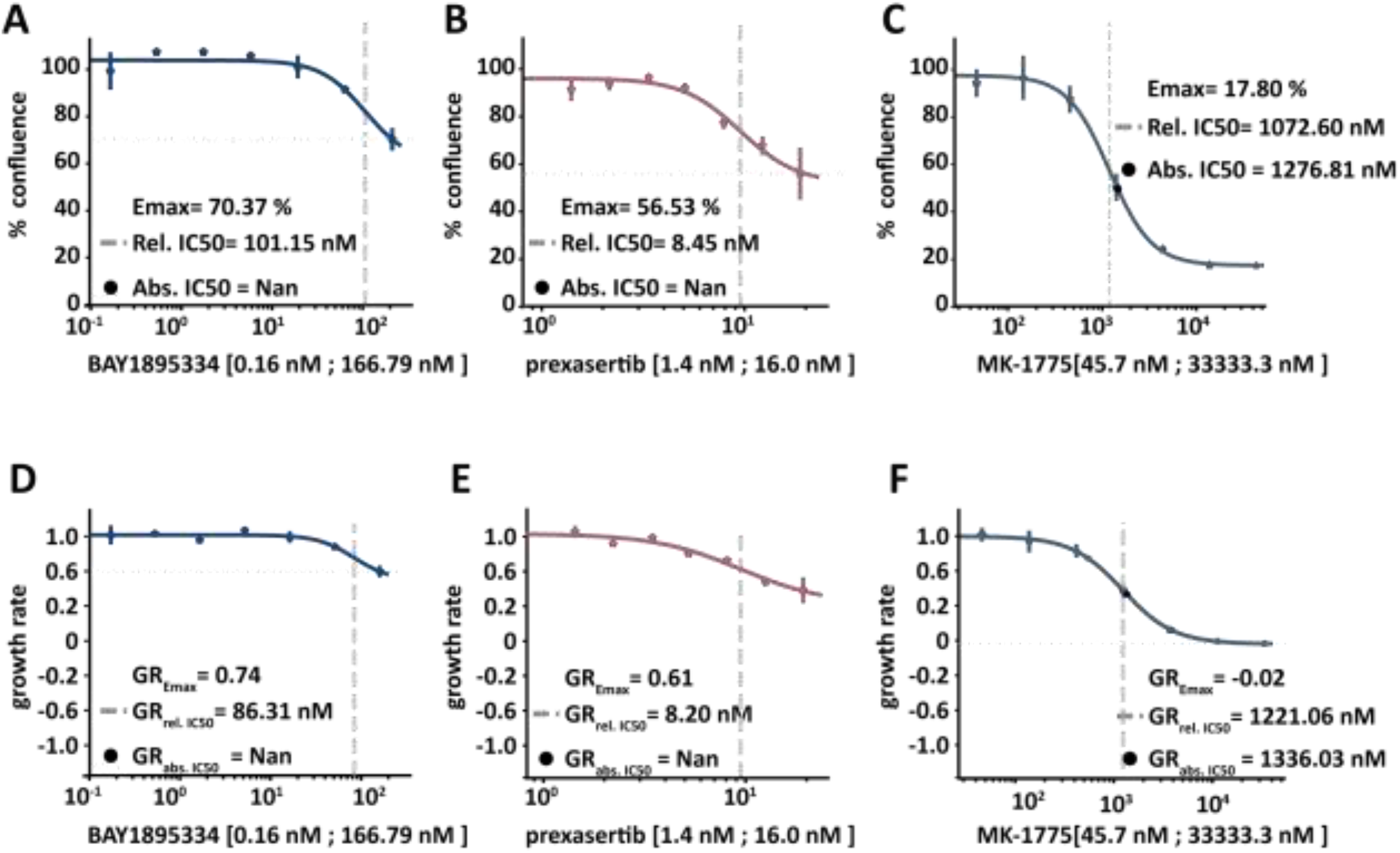
Dose-response curve provided by HTSplotter for BAY1895334, prexasertib and MK-1775 at 72h. **A)** Dose-response curve of BAY1895334, in which the Emax is 70.37% of confluency, the relative (rel.) IC50 is 101.15 nM and the absolute (abs.) IC50 could not be determined. **B)** Dose-response curve of prexasertib, in which the Emax is 56.53% of confluency, the rel. IC50 is 8.45 nM and the abs. IC50 could not be determined. **C)** Dose-response curve of MK-1775, in which the Emax is 17.80% of confluency, the rel. IC50 is 1072.60 nM and the abs. IC50 is 1276.81 nM. **D)** GR dose-response curve of BAY1895334, in which the GR_Emax_ is 0.74, the GR_rel. 50_ is 86.32 nM and the GR_abso. 50_ could not be determined. **E)** GR dose-response curve of prexasertib, in which the GR_Emax_ is 0.61, the GR_rel. 50_ is 8.20 nM and the GR_abs. 50_ could not be determined. **F)** GR dose-response curve of MK-1775, in which the GR_Emax_ is 0.02, the GR_rel. 50_ is 1221.05 nM and the GR_abs. 50_ is 1336.03 nM.

**Fig 7:**
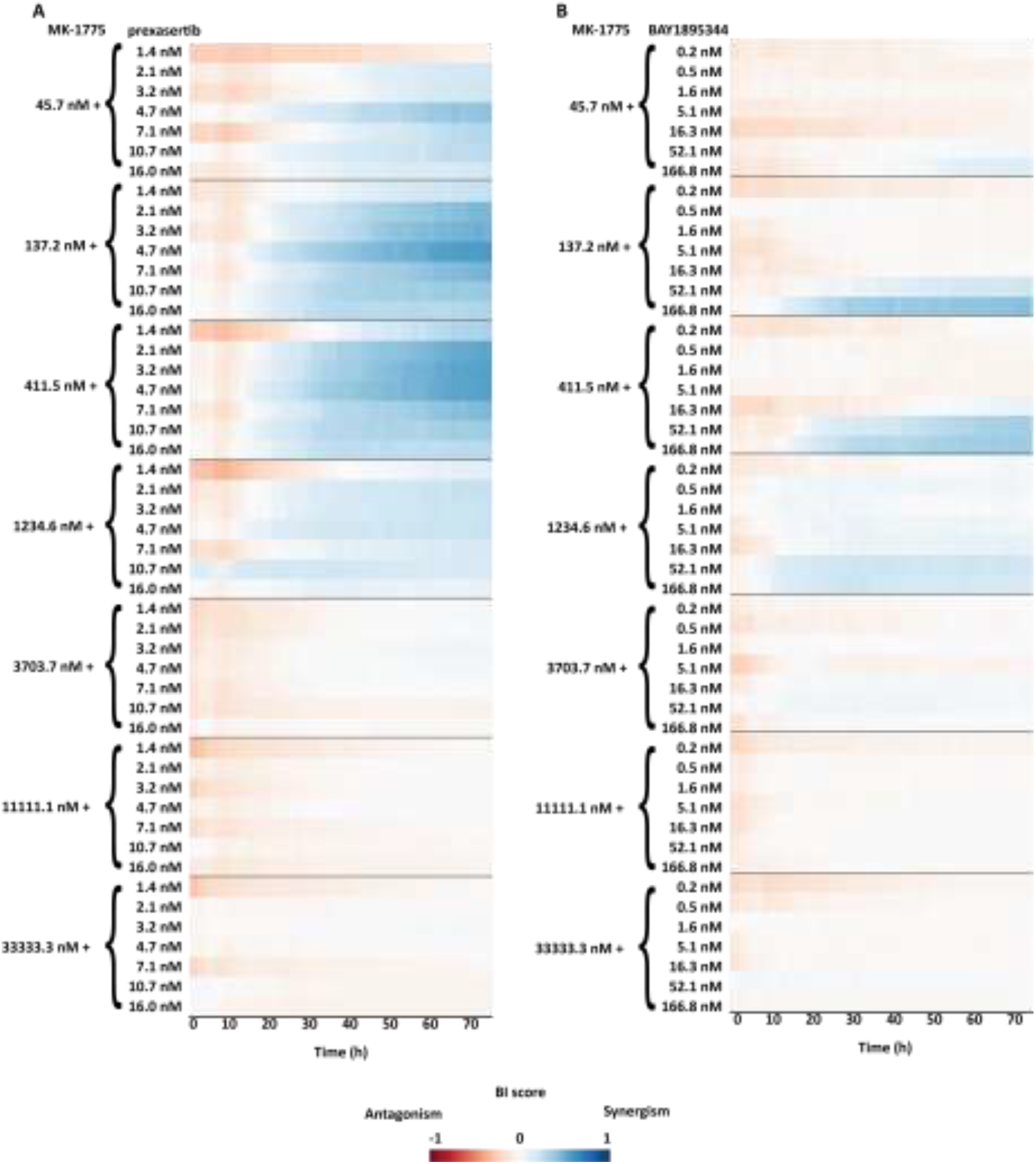
BI score heatmap of dose-effect combination overtime, in which the blue gradient indicates synergism and the red gradient indicates the antagonism. **A)** Dose-effect combination heatmap over time of a combination range of MK-1775 with a range of prexasertib (7 × 7 matrix). **B)** Dose-effect combination heatmap over time of a combination range of MK-1775 with range of BAY1895344 (7 × 7 matrix).

**Fig 8:**
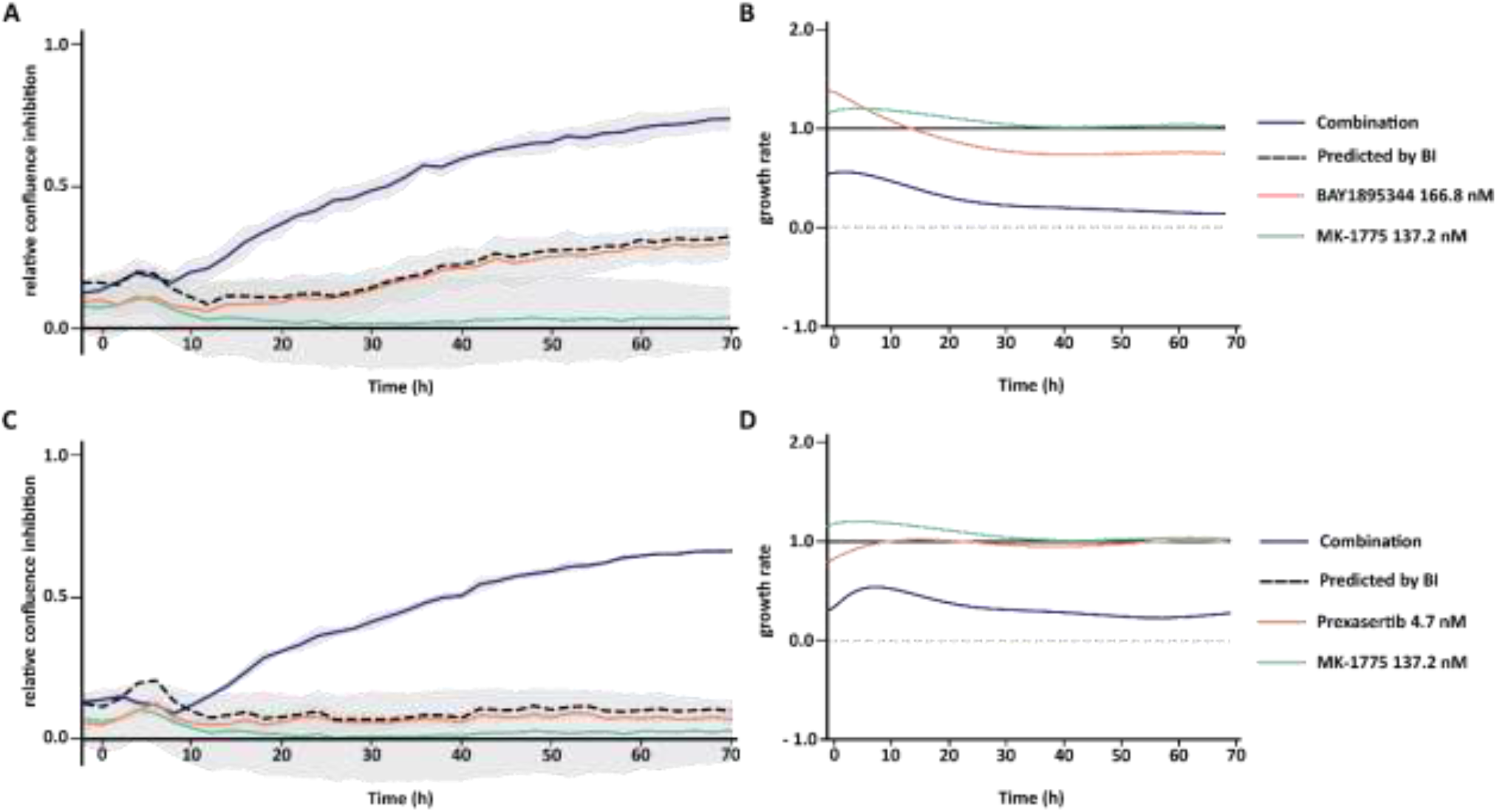
XY-plot and growth rate plot of the best dose-effect combination of MK-1775 with BAY1895344 (A, B) or MK-1775 with Prexasertib (C, D). Relative confluence is the confluence relative to the control. The dash line indicates the predicted combination effect, computed according to the BI method, equation (4). The growth rate is computed according to equation (7). The grey dash line on the growth rate plot indicates the stop on the growth rate. **A)** Relative confluence inhibition over time of BAY1895344 at 166.8 nM combined with MK-1775 at 137.2 nM. The BI scores at 24h, 48h and 72h are 0.28, 0.39 and 0.42, respectively. **B)** Growth rate of each condition in relation to the control. BAY1895344 growth rates at 24h, 48h and 72h, are 0.86, 0.74 and 0.74 respectively. MK-1775 growth rates at 24h, 48h and 72h, are 1.10, 1.02 and 1.01 respectively. The combined condition growth rates at 24h, 48h and 72h are 0.28, 0.18 and 0.14. **C)** Relative confluence inhibition plot over time of prexasertib at 4.7 nM combined with MK-1775 at 137.2 nM. The BI scores at 24h, 48h and 72h are 0.25, 0.47 and 0.56, respectively. **D)** In relation to the control. Prexasertib growth rates at 24h, 48h and 72h, are 1.00, 0.96 and 0.96 respectively. MK-1775 growth rates at 24h, 48h and 72h, are 1.10, 1.02 and 1.01 respectively. The combined condition growth rates at 24h, 48h and 72h are 0.37, 0.26, 0.29.

To illustrate the advantage of HTSplotter on drug combination evaluation assessed in real-time, we applied our tool to an in-house assessment of the human breast cancer cell line MCF-7 in response to single and combination exposure to three inhibitors targeting WEE1, CHK1 and ATR (MK-1775, prexasertib and BAY1895344, respectively). The combination of these inhibitors has been shown to be synergistic in different cancer entities, such as breast cancer (27)(28).. The drug combination experiment consisted of 136 conditions and 37 time points, resulting in 5032 data points. The data used in HTSplotter was directly exported from Incucyte S3.

Each drug alone was assessed by a dose-response, which provided the conventional metrics (abs. IC50, rel. IC50) as well as metrics based on the normalized growth rate inhibition (GR_abs 50_, GR_rel 50_ and GR_Emax_) (Fig 6). At 72h, the tested dose-range of MK-1775 allowed to determine the absolute IC50 and GR_abs 50_ (1276.21 nM, 1336.03 nM, respectively), as opposed to other drugs. Accordingly, the Emax and GR_Emax_ of both drugs, BAY1895334 and prexasertib, was above 50% of confluency and 0.5 of growth rate, confirming that either absolute IC50 and GR_abs 50_ were not achieved by the tested dose-range. Prexasertib showed the lowest relative IC50 and GR_rel 50_, while MK-1775 showed the highest relative IC50 and GR_rel 50_. Thus, prexasertib revealed to be the most potent drug, followed by BAY1895344 and MK-1775. As mentioned above, from the genetic-chemical perturbagen experiment analysis, the drug combination analysis also shows a more stable values from the GR metrics between the 24h, 48h and 72h, than the conventional ones (S Fig 5).

The drug combination setup encompassed a dose-response combination matrix of MK-1775 with BAY1895344 or prexasertib, referred as dose-effect combinations. The ZIP (S Fig 7) and BI (Fig 7) method were applied in order to evaluate synergism/antagonism.

HTSplotter revealed that prexasertib showed stronger synergism than BAY1895344 over time. For both combinations the BI heatmaps (Fig 7) show that the synergistic effects emerged at 20h of treatment for all combinations, while the ZIP heatmaps display that the synergistic effect started after 36h of treatment (S Fig 7). Moreover, for some synergistic combinations evaluated by both methods, the scores increased over time indicating an increasing synergistic effect in this cell line, unlike the other synergistic combinations. Increased BI and ZIP scores were less pronounced. Assessing in detail the highest BI score of each dose-effect combination (Fig 8, A and DError! Reference source not found.), BAY1895344 alone had a higher and increasing inhibitory effect when compared to MK-1775 and prexasertib, which both had almost no impact on the growth inhibition over time. Consequently, the predicted combination effect of MK-1775 with BAY1895344 increased over time, while the MK-1775 with prexasertib had a small and constant inhibition effect. Looking at the growth rate of each condition in relation to the control results (Fig 8, B and C), provided by HTSplotter, BAY1895344 and MK-1775 had a higher, but declining, growth rate, until 20h and 30h after treatment, respectively. After 40h of treatment, MK-1775 growth rate started to increase in relation to the control. Initially, the prexasertib condition alone had a lower but increasing growth rate, achieving the same level as the control over time. As for the combined condition, BAY1895344 with MK-1775, had an immediate decrease on the growth rate until 30h of treatment, stabilizing it over time. For the combined condition, prexasertib with MK-1775, the growth rate increased until 10h after treatment, and afterwards started to decrease until 60h after treatment. However, after the 60h time point the growth rate increases. Interestingly, this effect was not observed by the inhibition and BI results. Notably, assessing both analyses together, while the inhibition effect showed a stable synergistic effect, the complementary growth rate analysis suggests that these cells were still able to proliferate at a constant rate. Moreover, from the combination of MK-1775 with prexasertib, the increasing growth rate at the latest time point may raise the question if the identified turnover point from the growth rate is an indication of recovery phenotype from this combination. The recovery phenotype leads to the question if this is a resistance phenotype or perhaps a depletion of the drug, by intracellular mechanisms in the cell, which allowed these cells to increase proliferation rate again. In order to clarify these questions more biological replicates, as well as a more in depth analysis including transcriptomics and/or proteomics profiles are required. In case of a drug combination experiment, we showed that HTSplotter is efficient in processing and analysing data, providing the researcher important visualizations to facilitate data interpretation.

### Comparison to related software

Over the years, several tools were developed for the analysis of different types of HTS, such as drug combination and genetic-chemical perturbagen screens. Some of those, such as RNAither, IncucyteDRC and DRC, require programming knowledge (14)(29)(30) while others are more user friendly tools (Table 1) (13)(15)(22)(31)(32).

**Table 1.**
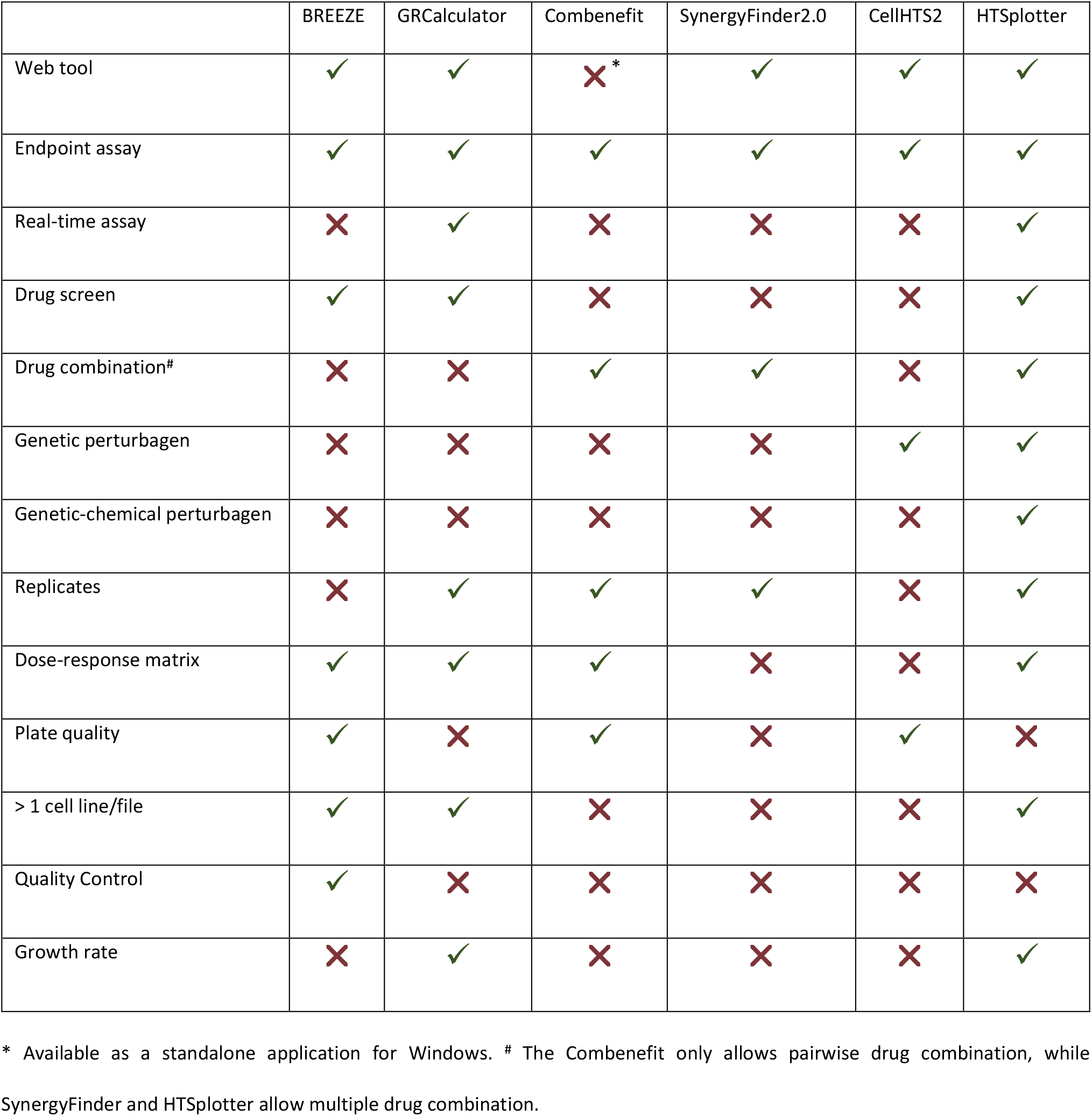
Open source software for HTS data analysis and their characteristics.

The available web tools and software are designed for specific experiment types, mostly endpoint assays. Often, one has to organize, structure and clean the data specifically to the tool. Depending on the selected tool, the user may have to normalize the data. Once the tool has been selected, this procedure has to be repeated for each selected time point. Therefore, for real-time assays at least the last step must be repeated as many times as selected time points. Additionally, in case of drug combination, some tools only provide a heatmap with synergism/antagonism scores, missing important information regarding the perturbation effect. For the genetic-chemical perturbagen screens, either an endpoint assay or an time course assay, there is no tool able to perform data analysis, necessitating researchers to conduct their data analysis manually.

To compare HTSplotter on drug combination with the available tools, we resorted to SynergyFinder, a tool widely used for synergism/antagonism evaluation. SynergyFinder was applied to the dose-effect combination at 72h using the BI method. The tailoring of the data to the specific input format of SynergyFinder was performed by HTSplotter. The dose-response curve profile matched the one provided by HTSplotter, however, the statistical metrics, such as χ^2^ (Chi-square) and R^2^ (R-squared), of the dose-response relationship of the single agent were not provided by SynergyFinder.

SynergyFinder identified the same synergistic dose range as the HTSplotter using the BI and ZIP methods (Fig 9, and S Fig 7), with minor differences in scores due to different approaches. SynergyFinder computes the average over the full dose-response matrix, while HTSplotter computes synergism between tested concentrations. Moreover, HTSplotter comes with many other features. When multiple time points are assessed, from a single analysis, HTSplotter provides a complete overview from the experiment results to the user by showing the synergism score heatmap over time, indicating if and when synergism starts and ends. Complementary to the heatmaps are the XY-plots and growth rate plots, which provide a more detailed analysis. These plots allow a comparison, in terms of growth rate and inhibition effect, between combination and each condition alone. Based on the growth rate profile over time upon a certain perturbation, these analyses allow the researcher to identify possible phenotypes such as drug resistance, senescence or cell death induction. In case of a unique time point experiment assessed by HTSplotter, a bar plot with the inhibition effect is also provided (S Fig 6**Error! Reference source not found**.). In case of combinations, either genetic-chemical or drug combinations, the researcher observes the effect of each drug alone, as well as the predicted and the observed effects upon combination, which may aid the optimization of further experiments in addition to the identification of the best concentrations for combination experiments. In contrast to other tools, if any experiment type consists of a time course assay, HTSplotter analysis the growth rate which may lead the user to the identification of different cellular phenotypes, such as cell death or senescence.

**Fig 9:**
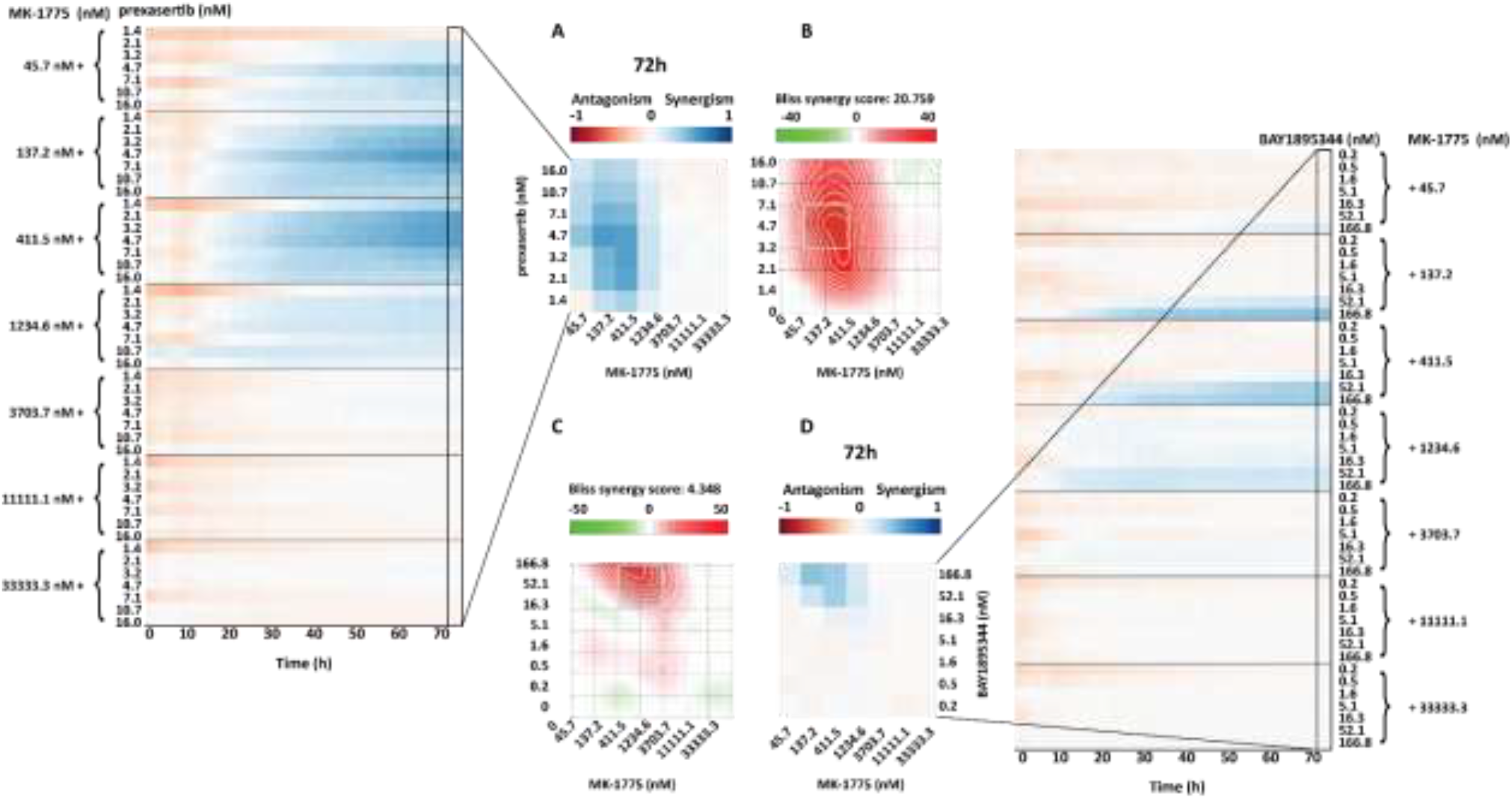
SynergyFinder 2.0 and HTSplotter heatmap at 72h. **A)** HTSplotter heatmap over time and at final time point (72h) of dose-effect combination of prexasertib with MK-1775, maximum BI score of 0.56 at 72h. **B)** SynergyFinder heatmap of dose-effect combination of prexasertib with MK-1775, with BI score of 20.759. **C)** SynergyFinder heatmap of dose-effect combination of BAY1895344 with MK-1775, with BI score of 4.348. **D)** HTSplotter heatmap over time and at final time point (72h) of dose-effect combination of BAY1895344 with MK-1775, maximum BI score of 0.42 at 72h. HTSplotter has a fixed legend scale from -1 to 1.

## Conclusion

Academic research is moving towards HTS, from dozens to thousands screenings per day, in order to be more efficient, to have more results and to answer faster biological questions. Additionally, the research in question might request not only a drug screen experiment but also genetic or genetic-chemical perturbagen screens. This implies that one researcher or a research group, have different types of data to be analysed in the shortest time possible. These results determine not only the next research steps but also experiment and research conclusions. Aiming to accelerate the research, several tools have been developed for the HTS data analysis. However, these lack analytical flexibility on the processing and downstream analyses for several time points and for genetic-chemical perturbagen screens. Although these tools already reduce the research time, the lack of standardization significantly augments the time preparing different input files and the data processing, overall delaying scientific progress. Of note, to the best of our knowledge, there is currently no tool available to perform the analysis of different types of experiments.

In order to address these research needs and to bring standardization across different experiment types, we developed HTSplotter which serves as a unique tool to analyse either drug, drug combination, genetic perturbagen and genetic-chemical perturbagen screens, both in endpoint and in real-time assays. Here, we demonstrated that HTSplotter enables a tailored end-to-end analysis, including processing, dose-response, synergism/antagonism and growth rate analyses, as well as high quality visualisations, requiring minimum user intervention. We also demonstrated that HTSplotter is fit for genetic-chemical combination screens, thus amenable for chemical-genetics (*e*.*g*. CRISPR/drug) experiments. Additionally, this tool also analyses higher-order drug combinations, combinations with more than two drugs. HTSplotter provides a unique, detailed visualisation of the drug combination effect and growth rate over time, going beyond the state-of-the-art, for a better biological interpretation. Hence, we believe that HTSplotter is a major contribution to biomedical research with efficient and effective data analysis providing results in a throughput manner for different type screenings. Future HTSplotter updates will focus on adding quality control process to the data analysis, incorporation of other dose-response curve models, such as bi-modal, and the integration of other synergistic/antagonistic models, such as HAND and BRAID.

## Materials and methods

### Dose-response curve and drug combination analysis

HTSplotter implemented a four-parameter logistic function, testing the curve fitness using the Levenberg-Marquardt algorithm for nonlinear least squares curve-fitting (SciPy 1.6.0). The curve fitting statistical parameters, such as Chi-square, residuals and R-squared were calculated. For the calculation of the AUC, the trapezoidal rule was applied (NumPy 1.19.5). To compute the different ICs, HTSplotter determined the four-parameter logistic function coefficients.

HTSplotter applies the BI, HAS and ZIP methods to calculate synergism/antagonism scores for pairwise or higher order combinations. Before computing the BI, HAS score for each time point, HTSplotter verified for each condition if the cell viability effect was higher than the control. If so, the effect was corrected to the maximum of the control, to avoid false BI and HSA scores.

In the case of a drug combination or genetic-chemical perturbagen screen, the method chosen by the user is applied by the HTSplotter in order to determine synergism/antagonism, from a combination of *N* drugs, in which the drugs *B*_1_, *B*_2_, …, *B*_*N*_ at doses *δ*_1_, *δ*_2_, … *δ*_*N*_, being *D*_1_, *D*_2_, …, *D*_*N*_, the inhibition effects of drugs *B*_1_, *B*_2_, …, *B*_*N*_ at doses *δ*_1_, *δ*_2_, … *δ*_*N*_, respectively. In order to compute the HSA score, first the highest inhibition effect is identified, equation (1) (18), and then the HSA score is obtained through the difference between the observed inhibition effect, 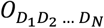 of the combination and HSA value, equation (2). As for the BI method, first HTSplotter determines the predicted combination effect by the BI method (20), equation (3), where *H(F)* is the predicted inhibition effect from the combination. Next, the BI score is determined by the difference between 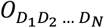 and *H(F)*, equation (4). Finally, and only for drug combination, HTSplotter determines the ZIP score, by the following the method described in (17), extended to higher orders according to equation (5).

If the synergism score is higher than zero, the combination is synergistic, if below zero then it is antagonistic. Following these analyses, data is exported as text and results are plotted in a PDF file.

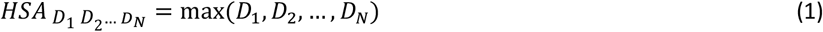

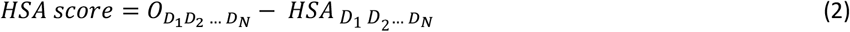

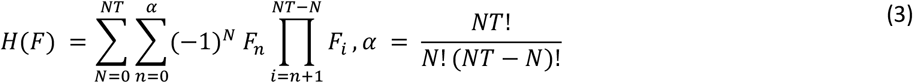

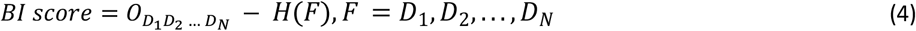

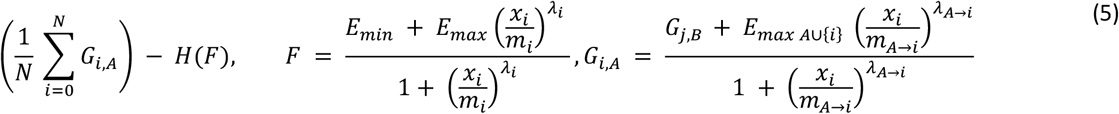

For the comparison study, SynergyFinder 2.0 was applied following their guidelines(32).

In order to determine the growth rate over time, HTSplotter first rescales each condition to the first time point and then derives the confluency. Then, HTSplotter implemented splines with minimum squares and differentiated using central finite differences, making use of two methods (based on fixed intervals and time dependent), relating them in order to construct smoother growth rate evolutions (21).

Considering three instances in time, *t*_1_ < *t* < *t*_2_, (S Fig 1) we can obtain the fixed intervals GR from *t* = 0 up to any of the time points, ⟨*GR*⟩(*c, t*), which can be seen as a mean GR in a set interval of time with condition *c*. So that ⟨*GR*⟩(*c, t*) is in fact the mean of GR over time, *GR*(*c, t*), in the interval [0, *t*], then definition (6) must be verified.

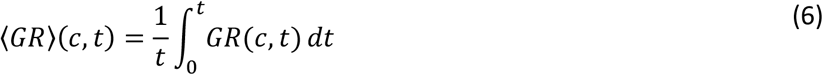

Using a Taylor series expansion, ⟨*GR*⟩(*c, t*) at time points *t*_1_and *t*_2_, centered at *t*, ignoring higher order terms, considering the definition (6) and making *t*_2_, *t*_1_ → *t*, then *GR*(*c, t*) can be written as in equation (7).

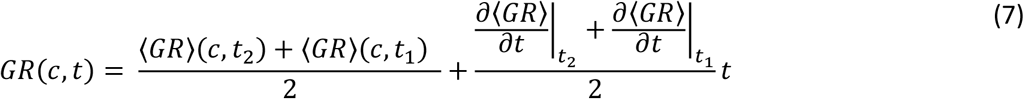

### Case study

For the case study, MCF-7 and SOX11 overexpression SH-EP cell line were grown in RPMI medium supplemented with 10% FCS and 2mM L-Glutamine and 100 IU/mL penicillin/streptomycin. The generation of SOX11 overexpression is described in (23). To evaluate synergism and the dose-response relationship, cells were seeded in a 384-well plate (Corning COS3764), at a density of 3×10^3^ cells per well. Cells were allowed to adhere overnight, after which these were exposed to the respective treatment.. The drugs used were: AZD2281 (Selleckchem, S1060), MK-1775 (Selleckchem, S1525), prexasertib (MedChem Express, HY-18174A) and BAY 1895344 (Selleckchem, S8666). The treatment was performed by the D300 TECAN. Cell proliferation was monitored for 72h, in which pictures were taken through IncuCyte S3 Live Cell Imaging System each 2h.Each image was analysed through the IncuCyte S3 Software. Cell proliferation was monitored by analysing the occupied area (% confluence) over time.

## Supporting information

**S Fig 1:**
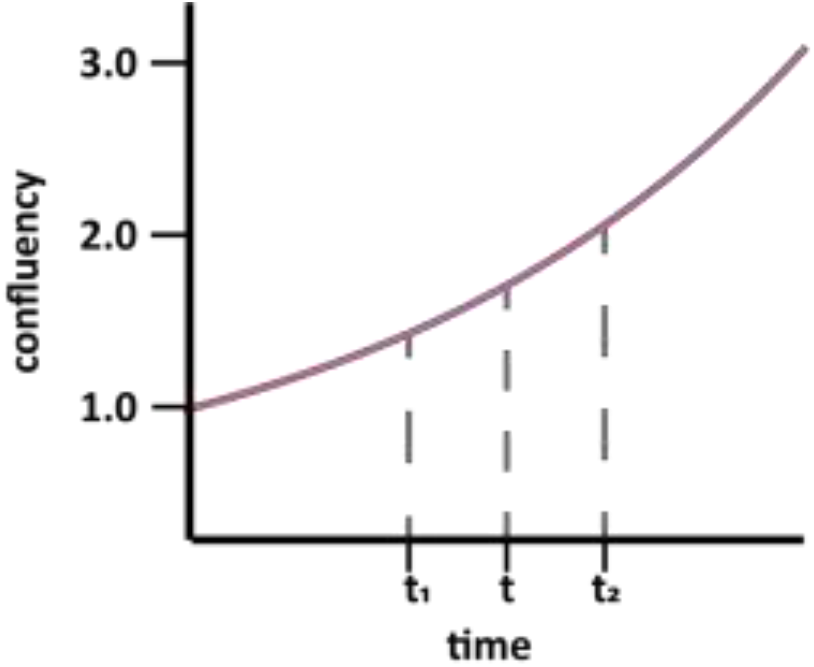
Time dependent value based on confluency over time.

**S Fig 2:**
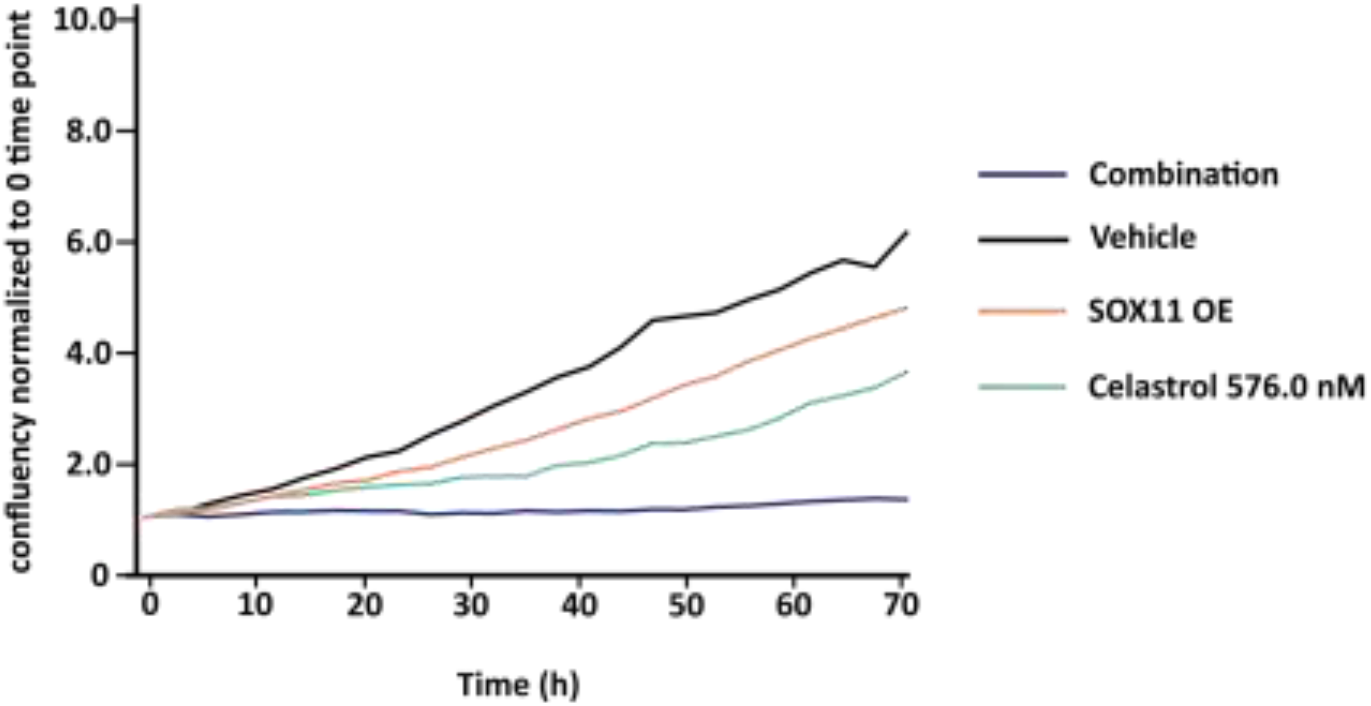
Fig. XY-plot normalized to the 0 time point provided by HTSplotter for the SOX11 OE and Celastrol. Confluency of each condition normalized to the confluence at the 0 time point. In contrast to the combined condition, each condition alone has an increase confluency over time.

**S Fig 3:**
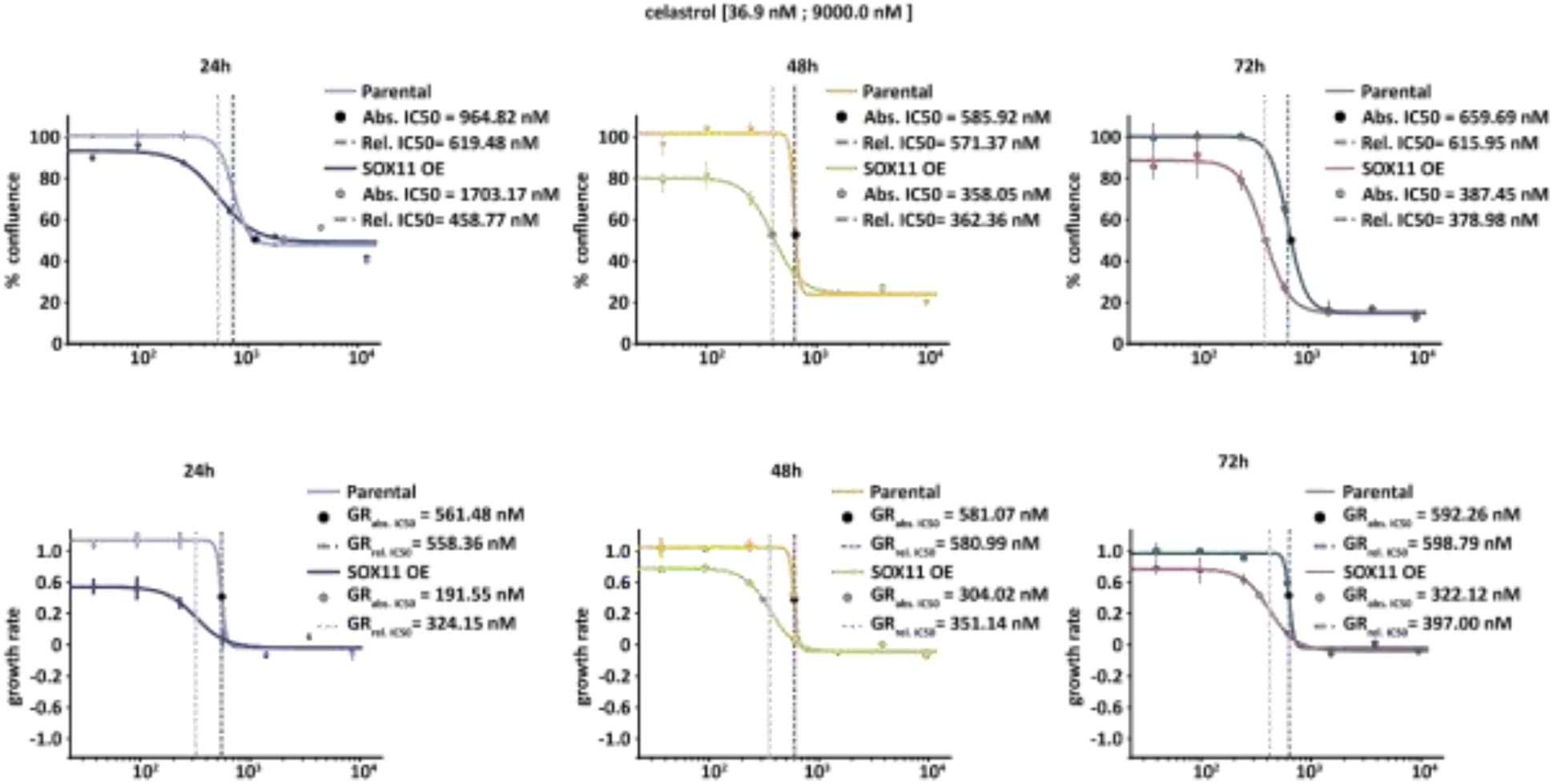
Celastrol dose-response curve analysis, where conventional and GR metrics are shown over time.

**S Fig 4:**
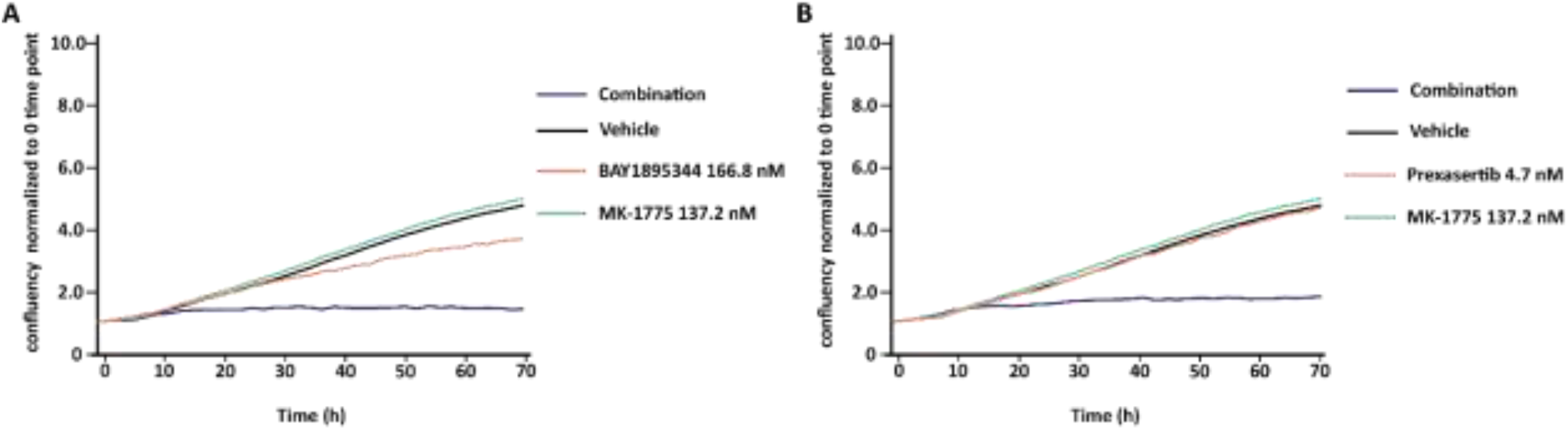
XY-plot normalized to 0 time point provided by HTSplotter, where the confluency of each condition is normalized to the confluence at the 0 time point. **A)** An increasing confluency is observed on the BAY1895344 and MK-1775 conditions alone. Over time MK-1775 have a higher confluency, when compared to the control. As the BAY1895344 condition the confluency is lower over time. As for the combined after 10h of treatment the confluency stabilized. **B)** An increasing confluency is observed for the prexasertib and MK-1775 conditions alone, being the MK-1775 slightly above from the control. The combined condition after 10h of treatment has a constant confluence.

**S Fig 5:**
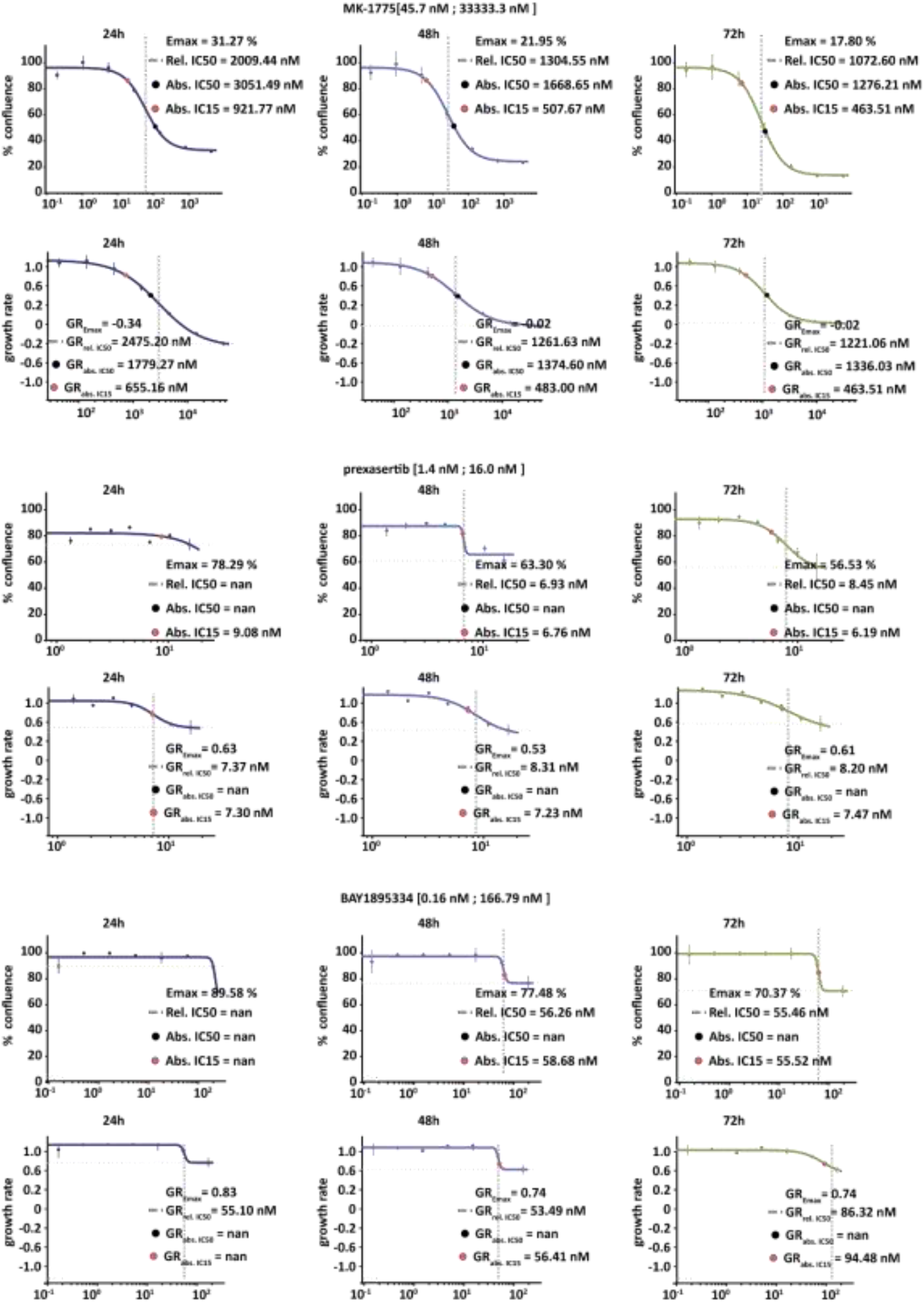
MK-1775, prexasertib and BAY1895344 dose-response curve analyses, where conventional and GR metrics are shown over time.

**S Fig 6:**
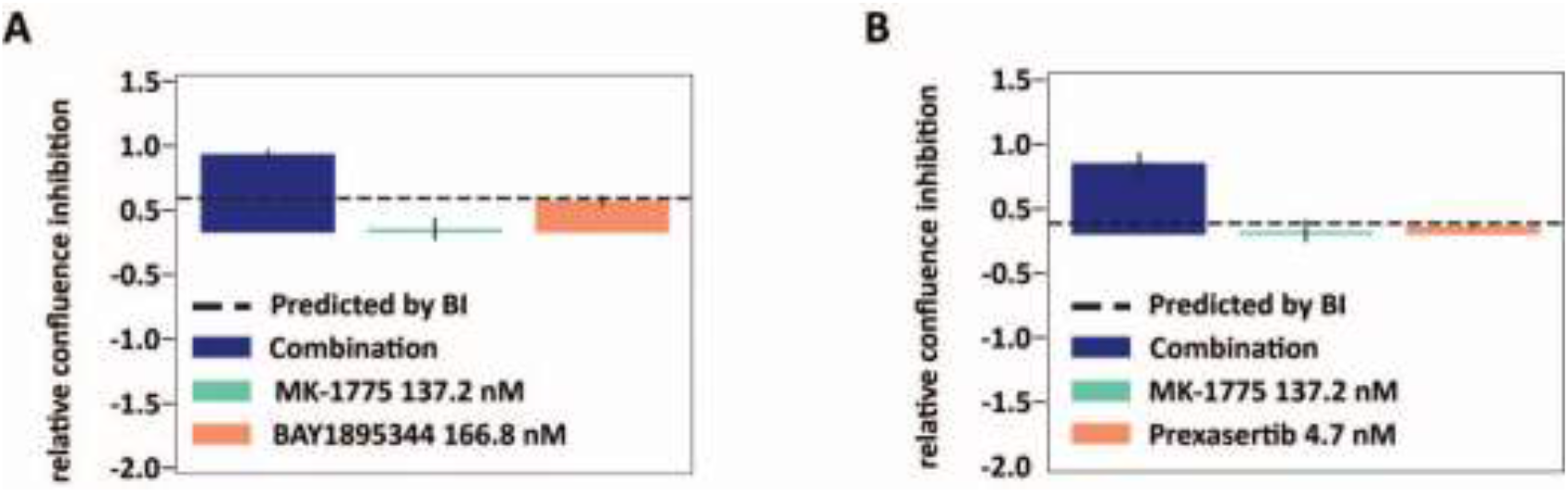
Bar plot, at 72h, of the best dose-effect combination of MK-1775 with BAY1895344 (A) or MK-1775 with Prexasertib (B). The dash line indicates the predicted combination effect, computed according to the BI method, equation (3). **A)** Relative confluence inhibition of BAY1895344 at 166.8 nM combined with MK-1775 at 137.2 nM, with a BI score of 0.42. **B)** Relative confluence inhibition of prexasertib at 4.7 nM combined with MK-1775 at 137.2 nM, with BI a score of 0.56.

**S Fig 7:**
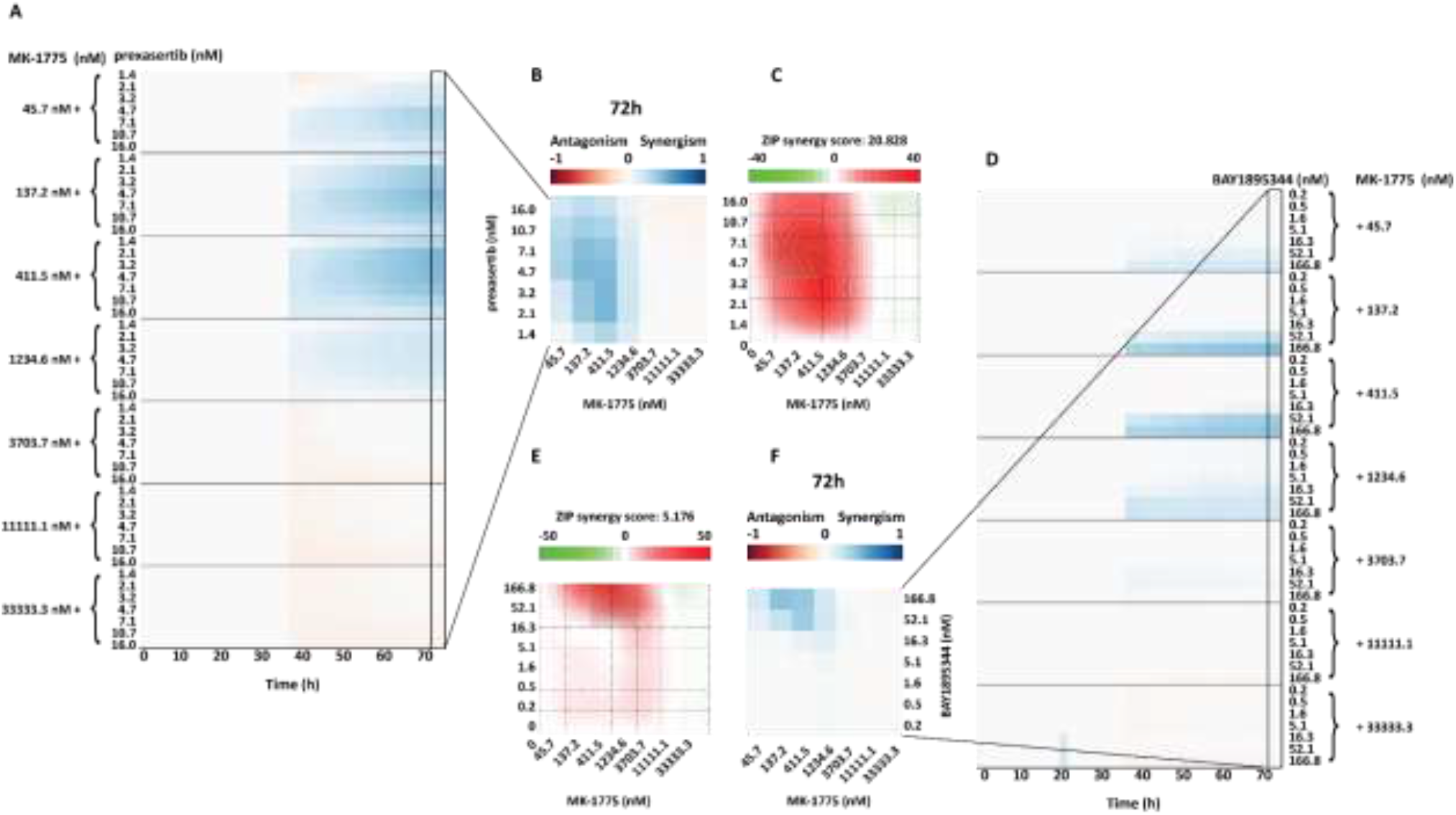
SynergyFinder 2.0 and HTSplotter heatmap at 72h. **A)** HTSplotter heatmap over time and **B)** at final time point (72h) of dose-effect combination of prexasertib with MK-1775, maximum ZIP score of 0.46 at 72h. **C)** SynergyFinder heatmap of dose-effect combination of prexasertib with MK-1775, with ZIP score of 20.828. **D)** HTSplotter heatmap over time and **E)** at final time point (72h) of dose-effect combination of BAY1895344 with MK-1775, maximum ZIP score of 0.41 at 72h. **F)** SynergyFinder heatmap of dose-effect combination of BAY1895344 with MK-1775, with ZIP score of 5.176. HTSplotter has a fixed legend scale from -1 to 1.

## Acknowledgments

The authors acknowledge the following individuals for providing data from their projects: Eric de Bony de Lavergne, Jolien Van Laere, Siebe Loontiens and Suzanne Vanhauwaert.

## Author contributions

**Conceptualization** - Jolien De Wyn, Carolina Nunes, Frank Speleman, Jo Vandesompele and Vanessa Vermeirssen

**Formal Analysis** - Carolina Nunes

**Investigation** - Fanny De Vloed and Carolina Nunes

**Methodology** - Carolina Nunes

**Software** - Jasper Anckaert and Carolina Nunes

**Supervision** - Frank Speleman and Vanessa Vermeirssen

**Visualisation** - Carolina Nunes and Vanessa Vermeirssen

**Writing - original draft** - Carolina Nunes and Vanessa Vermeirssen

**Writing - review & editing** - Kaat Durinck, Carolina Nunes, Frank Speleman, Jo Vandesompele, and Vanessa Vermeirssen

## References

1. Falzone L, Salomone S, Libra M. Evolution of cancer pharmacological treatments at the turn of the third millennium. Front Pharmacol. 2018;9(NOV).

2. Mott BT, Eastman RT, Guha R, Sherlach KS, Siriwardana A, Shinn P, et al. High-throughput matrix screening identifies synergistic and antagonistic antimalarial drug combinations. Sci Rep. 2015;5(August):1–14.

3. Jeon M, Kim S, Park S, Lee H, Kang J. In silico drug combination discovery for personalized cancer therapy. BMC Syst Biol. 2018;12(Suppl 2).

4. Rothan HA, Teoh TC. Cell-Based High-Throughput Screening Protocol for Discovering Antiviral Inhibitors Against SARS-COV-2 Main Protease (3CLpro). Mol Biotechnol [Internet]. 2021;63(3):240–8. Available from: https://doi.org/10.1007/s12033-021-00299-7

5. Meyer CT, Wooten DJ, Lopez CF, Quaranta V. Charting the Fragmented Landscape of Drug Synergy. Trends Pharmacol Sci [Internet]. 2020;41(4):266–80. Available from: https://doi.org/10.1016/j.tips.2020.01.011

6. Inglese J, Johnson RL, Simeonov A, Xia M, Zheng W, Austin CP, et al. High-throughput screening assays for the identification of chemical probes. Nat Chem Biol. 2007;3(8):466–79.

7. Sham O, Sanjana NE, Zhang F. High-throughput functional genomics using CRISPR-Cas9. Nat Rev Genet. 2015;16(5):299–311.

8. Sun X, Vilar S, Tatonetti NP. High-throughput methods for combinatorial drug discovery. Sci Transl Med. 2013;5(205).

9. Williams SP, Barthorpe AS, Lightfoot H, Garnett MJ, McDermott U. Data Descriptor: High-throughput RNAi screen for essential genes and drug synergistic combinations in colorectal cancer. Sci Data. 2017;4:1–13.

10. Lehár J, Stockwell BR, Giaever G, Nislow C. Combination chemical genetics. Nat Chem Biol. 2008;4(11):674–81.

11. Single A, Beetham H, Telford BJ, Guilford P, Chen A. A comparison of real-time and endpoint cell viability assays for improved synthetic lethal drug validation. J Biomol Screen. 2015;20(10):1286–93.

12. Ianevski A, He L, Aittokallio T, Tang J. SynergyFinder: a web application for analyzing drug combination dose–response matrix data. Stegle O, editor. Bioinformatics [Internet]. 2017 Aug 1;33(15):2413–5. Available from: https://academic.oup.com/bioinformatics/article/33/15/2413/3100330

13. Di Veroli GY, Fornari C, Wang D, Mollard S, Bramhall JL, Richards FM, et al. Combenefit: An interactive platform for the analysis and visualization of drug combinations. Bioinformatics. 2016;32(18):2866–8.

14. Ritz C, Baty F, Streibig JC, Gerhard D. Dose-response analysis using R. PLoS One. 2015;10(12):1–13.

15. Clark NA, Hafner M, Kouril M, Williams EH, Muhlich JL, Pilarczyk M, et al. GRcalculator: an online tool for calculating and mining dose-response data. BMC Cancer. 2017;17(1):698.

16. Yadav B, Pemovska T, Szwajda A, Kulesskiy E, Kontro M, Karjalainen R, et al. Quantitative scoring of differential drug sensitivity for individually optimized anticancer therapies. Sci Rep. 2014;4:1–10.

17. Yadav B, Wennerberg K, Aittokallio T, Tang J. Searching for drug synergy in complex dose–response landscapes using an interaction potency model. Comput Struct Biotechnol J [Internet]. 2017;15:387. Available from: http://dx.doi.org/10.1016/j.csbj.2015.09.001

18. Foucquier J, Guedj M. Analysis of drug combinations: current methodological landscape. Pharmacol Res Perspect [Internet]. 2015;3(3):e00149. Available from: http://www.pubmedcentral.nih.gov/articlerender.fcgi?artid=4492765&tool=pmcentrez&rendertype=abstract%5Cn http://doi.wiley.com/10.1002/prp2.149

19. Sinzger M, Vanhoefer J, Loos C, Hasenauer J. Comparison of null models for combination drug therapy reveals Hand model as biochemically most plausible. Sci Rep [Internet]. 2019;9(1):1–16. Available from: http://dx.doi.org/10.1038/s41598-019-38907-x

20. Zhao W, Sachsenmeier K, Zhang L, Sult E, Hollingsworth RE, Yang H. A new bliss independence model to analyze drug combination data. J Biomol Screen. 2014;19(5):817–21.

21. Hafner M, Niepel M, Chung M, Sorger PK. Growth rate inhibition metrics correct for confounders in measuring sensitivity to cancer drugs. Nat Methods [Internet]. 2016 Jun 2;13(6):521–7. Available from: http://www.nature.com/articles/nmeth.3853

22. Potdar S, Ianevski A, Mpindi JP, Bychkov D, Fiere C, Ianevski P, et al. Breeze: An integrated quality control and data analysis application for high-throughput drug screening. Bioinformatics. 2020;36(11):3602–4.

23. Decaesteker B, Louwagie A, Loontiens S, De Vloed F, Roels J, Vanhauwaert S, et al. SOX11 is a lineage-dependency factor and master epigenetic regulator in neuroblastoma. bioRxiv [Internet]. 2020;2020.08.21.261131. Available from: https://doi.org/10.1101/2020.08.21.261131

24. Uttarkar S, Frampton J, Klempnauer KH. Targeting the transcription factor Myb by small-molecule inhibitors. Exp Hematol [Internet]. 2017;47:31–5. Available from: http://dx.doi.org/10.1016/j.exphem.2016.12.003

25. Coulibaly A, Haas A, Steinmann S, Jakobs A, Schmidt TJ, Klempnauer K. The natural anti-tumor compound Celastrol targets a Myb-C/EBPβ-p300 transcriptional module implicated in myeloid gene expression. Bunting KD, editor. PLoS One [Internet]. 2018 Feb 2;13(2):e0190934. Available from: https://dx.plos.org/10.1371/journal.pone.0190934

26. Rajbhandari P, Lopez G, Capdevila C, Salvatori B, Yu J, Rodriguez-Barrueco R, et al. Cross-cohort analysis identifies a TEAD4–MYCN positive feedback loop as the core regulatory element of high-risk neuroblastoma. Cancer Discov. 2018;8(5):582–99.

27. Lewis CW, Jin Z, Macdonald D, Wei W, Qian XJ, Choi WS, et al. Prolonged mitotic arrest induced by Wee1 inhibition sensitizes breast cancer cells to paclitaxel. Oncotarget. 2017;8(43):73705–22.

28. Carrassa L, Chilà R, Lupi M, Ricci F, Celenza C, Mazzoletti M, et al. Combined inhibition of Chk1 and Wee1: In vitro synergistic effect translates to tumor growth inhibition in vivo. Cell Cycle. 2012;11(13):2507–17.

29. Rieber N, Knapp B, Eils R, Kaderali L. RNAither, an automated pipeline for the statistical analysis of high-throughput RNAi screens. Bioinformatics. 2009;25(5):678–9.

30. Chapman PJ, James DI, Watson AJ, Hopkins G V., Waddell ID, Ogilvie DJ. IncucyteDRC: An R package for the dose response analysis of live cell imaging data [version 1; referees: 2 approved]. F1000Research. 2016;5(May 2016):1–9.

31. Ianevski A, Giri AK, Aittokallio T. SynergyFinder 2.0: Visual analytics of multi-drug combination synergies. Nucleic Acids Res. 2021;48(1):W488–93.

32. Pelz O, Gilsdorf M, Boutros M. Web cellHTS2: A web-application for the analysis of high-throughput screening data. BMC Bioinformatics. 2010;11.

33. Roell KR, Reif DM, Motsinger-Reif AA. An introduction to terminology and methodology of chemical synergy-perspectives from across disciplines. Front Pharmacol. 2017;8(APR):1–11.

